# Pro-oxidant influence of quercetin supplementation in *Saccharomyces cerevisiae*

**DOI:** 10.1101/2024.06.10.598355

**Authors:** Andres Carrillo-Garmendia, Ana Leticia Vaca-Martinez, Blanca Lucia Carmona-Moreno, Juan Carlos González-Hernández, Jose Angel Granados-Arvizu, Sofia Maria Arvizu-Medrano, Jorge Gracida, Gerardo M. Nava, Carlos Regalado-Gonzalez, Luis Alberto Madrigal-Perez

**Affiliations:** Universidad Autónoma de Querétaro, Cerro de las Campanas, Santiago de Querétaro, Qro, 76010, México; Tecnológico Nacional de México/ ITS de Ciudad Hidalgo, Av. Ing. Carlos Rojas Gutiérrez #2120, Ciudad Hidalgo, Michoacán, 61100, México; Tecnológico Nacional de México/ IT de Morelia, Av. Tecnológico de Morelia, Morelia, Michoacán, 58030, México

**Keywords:** Quercetin, pro-oxidant, antioxidant systems, mitochondrial respiration, *Saccharomyces cerevisiae*, free radicals

## Abstract

How could quercetin exert a pro-survival phenotype (antioxidant) in mammalian cells while toxic to other eukaryotic cells? The redox capacity of quercetin may explain its antioxidant and toxic effects, based on the idea that quercetin impairs the electron transport chain, affecting ATP production and forming quercetin-derived free radicals. Herein, we provide evidence that quercetin supplementation: 1) depolarizes the mitochondrial membrane and augments the ADP/ATP ratio; 2) increases superoxide anion cellular levels; 3) changes the cellular response to H_2_O_2_ challenge associated with the antioxidant cellular response; 4) sensitizes the cellular response to lipoperoxidation challenge, and 5) augments cardiolipin levels. These events suggest that the quercetin pro-oxidant effect is related to mitochondrial respiration dysfunction and could induce cellular antioxidant response.

## Introduction

Quercetin (3,3’,4’,5,7□pentahydroxyflavone) is a flavonoid found in diverse fruits and vegetables, which presents a diversity of potential pharmacological applications, including a neuroprotective effect [1], cardiovascular protection [2], and anticancer properties [3]. Interestingly, quercetin has also shown toxicity to some organisms like *S. cerevisiae* [4], mammalian cell lines [5] and bacteria [6]. These apparently opposite phenotypes (beneficial and toxic) could be explained by quercetin interaction with the mitochondria [7], raising the hypothesis that quercetin could impair electron flux in the electron transport chain (ETC), producing a decrease in ATP generation. Furthermore, this idea also suggests that quercetin interaction with the ETC could generate reactive oxygen species (ROS) through quercetin-radical molecules or electron leakage [7]. Thus, it is possible that the antioxidant effect associated to quercetin could originate from the cellular antioxidant response and not from the molecule *per se*. Accordingly, quercetin augments nuclear translocation of the transcription factor Nrf2 that responds to oxidative stress and thioredoxin transcription, and activity in granuloma cells [8]. Besides, a report has shown that quercetin supplementation in HepG2 cells increases GSH content and induces transcription of the glutathione-peroxidase, glutathione-reductase, and glutathione-S-transferase genes [9]. Nonetheless, the molecular mechanism behind this pro-oxidant phenotype has not been completely explained.

Thus, this study aimed to identify whether the antioxidant phenotype exerted by quercetin is originated by a response to oxidative stress raised by mitochondria dysfunction. Herein, we report that quercetin supplementation affects mitochondrial membrane potential, ATP/ADP ratio, anion superoxide levels, and cellular growth under H_2_O_2_ and lipoperoxidation challenges.

## Material and methods

### Yeast strain and culture media

*S. cerevisiae* BY4742 (*Mat*α; *his3*Δ*1; leu2*Δ*0; lys2*Δ*0; ura3*Δ*0*) and strains gene-deletants *yap1*Δ, *ctt1*Δ, *sod2*Δ, *msn2*Δ and *gpx2*Δ were acquired from EUROSCARF (Frankfurt, Germany). YPD media (1% yeast extract, 2% peptone, 2% dextrose; BD Bioxon, Cuautitlan Izcalli, Mexico) were used for strain maintenance. The synthetic complete (SC) medium was also used for some experiments and consisted of 0.18% yeast nitrogen base without amino acids, 1% yeast synthetic drop-out supplements without uracil, and 400 μg/mL uracil (Merck, Darmstadt, Germany), 0.5% ammonium sulfate, 0.2% dipotassium phosphate (J.T. Baker, Phillipsburg, NJ, USA). In addition, the SC medium was supplemented with different concentrations of glucose and quercetin hydrate ≥95% (Merck). Geneticin (200 μg/mL; Merck) was added to the media for the deletant strains. Quercetin was dissolved in absolute ethanol (Sigma-Aldrich, St Louis MO, USA).

### Mitochondrial membrane potential

The mitochondrial membrane potential determination was performed employing the mitochondrial membrane potential kit (Merck; MAK159). The cells were grown in 5 mL SC medium supplemented with glucose (0.1%, 0.5%, or 5%) and quercetin (0.1 μM or 100 μM) at 30 °C and 200 rpm in an orbital shaker (Thermo Scientific, MaxQ 600, Marietta, OH, USA). Cells in the mid-log phase (OD_600_ ~ 0.5) were centrifuged at 4000 x *g* for 4 min, and the cell pellet was resuspended in PBS solution. A 96-well black microplate was added with 50 μL cell suspension (3 × 10^6^ cells/well) and followed the manufacturer’s instructions. Ratio fluorescence analysis evaluated the intensity of red fluorescence (excitation = 490, emission = 525 nm) and green fluorescence (excitation = 540, emission = 590 nm).

### ADP/ATP ratio

The ADP/ATP ratio was quantified employing the ADP/ATP ratio assay kit (Merck; MAK135). *S. cerevisiae* was grown in 5 mL SC medium supplemented with glucose (0.1%, 0.5%, 2%, or 5%) and quercetin (0.1 μM or 100 μM) at 30 °C and 200 rpm in an orbital shaker (Thermo Scientific, MaxQ™ 600). Cells were grown to mid-log phase (OD_600_ ~ 0.5) and 10 μL (2 × 10^3^ cells/mL) were transferred into a 96-well microplate. Subsequently, manufacturer specifications were followed to determine luminiscence (Varioskan Flash, Thermo Scientific; Vantaa, Finland).

### Reactive oxygen species analysis

Total ROS levels quantification was carried out using 2′,7′-dichlorofluorescein diacetate (H_2_DCFDA; Merck) assay. Cells were grown to mid-log phase (2 × 10^7^ CFU/mL, OD_600_ ~ 0.5), using SC media supplemented with glucose (0.1%, 0.5%, 2%, or 5%) and quercetin (0.1 μM or 100 μM) at 30 °C, mixed with H_2_DCFDA (5 mg/mL), and incubated for 1 h at 28 °C, with gentle shaking. Cells were washed twice with PBS solution, and fluorescence was measured at an excitation wavelength of 484 nm and 518 nm emission, employing a Varioskan Flash (Thermo Scientific).

The anion superoxide determination was performed using MitoSOX™ mitochondrial superoxide indicator (Thermo Scientific, Waltham, MA, USA), according to manufacturer instructions. *S. cerevisiae* cultures were grown to mid-log phase (OD_600_ ~ 0.5) in 5 mL SC medium supplemented with glucose (0.1%, 0.5%, 2%, or 5%) and quercetin (0.1 μM or 100 μM) at 30 °C, and 200 rpm in an orbital shaker (Thermo Scientific, MaxQ 600). For MitoSOX™ evaluation, the cells were placed in a black microplate (2 × 10^7^ cells/well), and fluorescence was measured at an excitation wavelength of 563 nm and an emission of 587 employing a Varioskan Flash (Thermo Scientific).

### H_2_O_2_ challenge

*S. cerevisiae* strains (WT, *msn2*Δ, *ctt1*Δ, *sod2*Δ, *yap1*Δ, and *gpx2*Δ) were grown in YPD media supplemented with two glucose concentrations (0.5% or 5%) and two quercetin levels (0.1 or 100 μM) at 30 °C with constant shaking. Cells were harvested in the mid-log phase (DO_600_ ~ 0.5) by centrifugation at 5000 x *g* for 3 min, washed twice with sterilized deionized water, and the cell pellet was resuspended in 400 μL sterilized deionized water. Then, 142 μL of YPD media, 3 μL of H_2_O_2_ (1 μM or 10 μM), and 5 μL of cell suspension were added into a 96-well microplate, incubated at 30 °C with constant shaking for 24 h, and the absorbance (600 nm) was recorded every hour employing a Multiskan Sky microplate reader (Thermo Scientific). The specific growth rate was calculated by fitting data with the exponential growth equation using the GraphPad Prism 10.1.1 software (La Jolla, CA, USA).

### Lipoperoxidation-inducing growth assays

The pre-inoculum of *S. cerevisiae* was grown overnight in 3 mL of YPD medium supplemented with 2% glucose with constant shaking at 30 °C. Subsequently, 10 μL of the cell suspension was added to 140 μL of SC medium supplemented with two levels of quercetin (0.1, or 100 μM) in combination with two glucose concentrations (0.1% or 5%), and placed into 96 wells microplate containing 100 μM of ferrous sulfate.

Another experimental design included three 1 mM fatty acids: palmitic 16:0, oleic 18:1(C9), and linoleic 18:2 (C9, C12); saturated, monounsaturated, and polyunsaturated, respectively, and a control without fatty acids. The microplate was incubated at 30 °C with constant shaking for 24 h, and absorbance (600 nm) was recorded every hour employing a Multiskan Sky microplate reader (Thermo Scientific). The specific growth rate was calculated by fitting data with the exponential growth equation using the GraphPad Prism 10.1.1 software (La Jolla, CA, USA).

### Cardiolipin quantification

Isolated mitochondria from the WT strain grown in SC medium supplemented with 0.1% or 5% glucose and quercetin (0.1, or 100 μM) at mid-log phase (DO_600_ ~ 0.5) were used for cardiolipin assay. Briefly, 50 μL of mitochondria (2.5 mg protein/mL) were used; 10 μM of acridine orange dye was added, and the volume was adjusted to 150 μl with 50 mM Tris/HCl buffer (pH 7.4), in a 96-well microplate. The samples were incubated for 10 min, then centrifuged at 12000 x *g* for 10 min. The concentration of acridine orange not bound to the membranes was calculated from a calibration curve (0-10 μM, at 495 nm), and the acridine orange bound to cardiolipin was calculated by subtracting the free from the total acridine orange. The cardiolipin concentration was calculated considering a stoichiometry of 2 moles of acridine orange/mol of cardiolipin.

### Statistical analysis

At least three independent experiments with two technical replicates were performed for the assays. Statistical analyses were computed using the GraphPad Prism 10.1.1 software (La Jolla, CA).

## Results

### Quercetin impact on mitochondrial membrane polarization

Quercetin impairs the electron flux of the ETC between ubiquinone pool and cytochrome *c* [4], which could promote electron leakage or quercetin-derived free radicals inducing oxidative stress. A key part of our hypothesis is the impairment of electron flux in the ETC by quercetin supplementation, which could cause mitochondrial membrane depolarization that was tested by measuring its membrane potential. Two glucose concentrations were used, aiming to exert different effects on mitochondrial respiration, 0.5% has shown higher response than 5% [10]. Additionally, two quercetin levels (0.1 μM and 100 μM) with dissimilar responses on mitochondrial respiration, and respiratory phenotypes were used [4]. As expected, *S. cerevisiae* quercetin supplementation decreased the mitochondrial membrane potential upon the three glucose concentrations tested (**Fig. 1a-c**). Notably, 100 μM quercetin showed a drastic decrease of the mitochondrial membrane potential compared to 0.1 μM quercetin supplementation at 0.5% and 5% glucose (*P* < 0.001) (**Fig. 1b-c**).

**Fig. 1.**
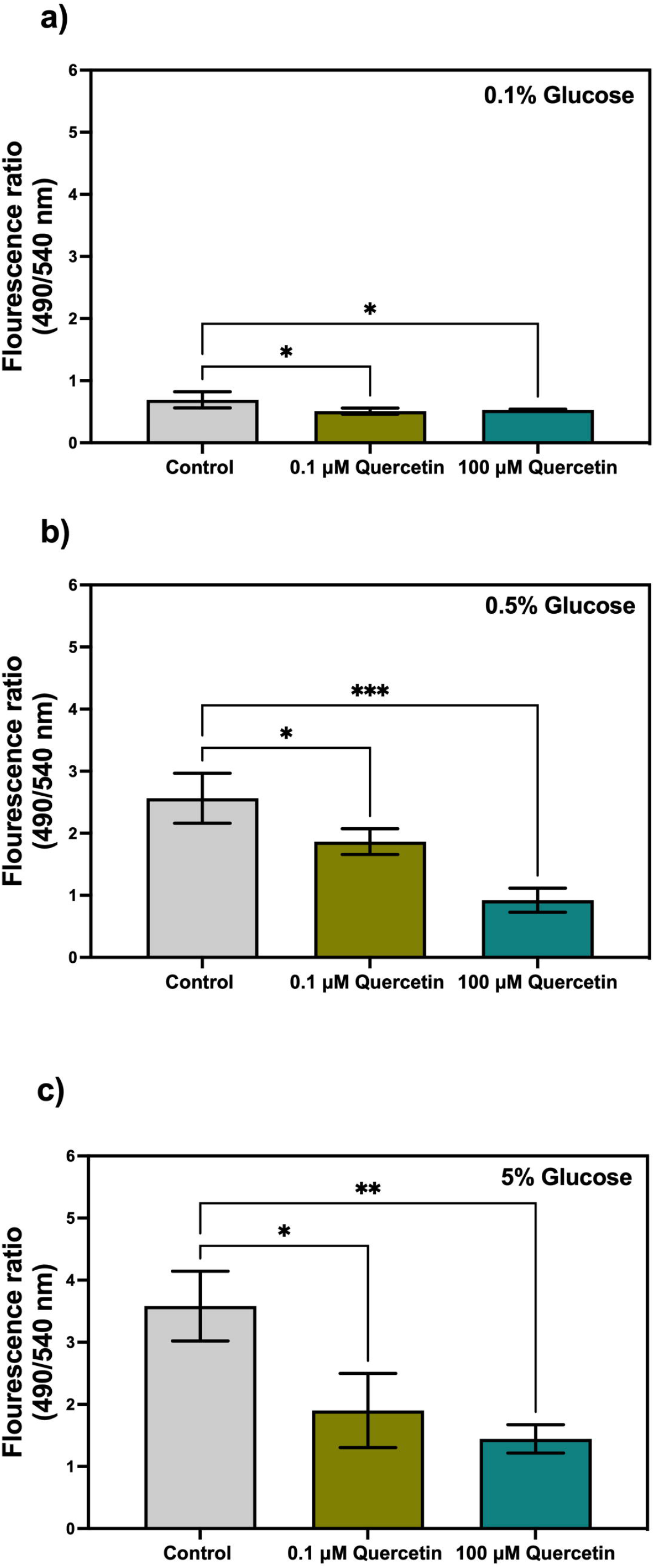
Influence of quercetin upon mitochondrial membrane potential of *S. cerevisiae*. Mitochondrial membrane potential was measured by fluorescence in mid-exponential cultures grown with SC media supplemented with **a)** 0.1% glucose: **b)** 0.5% glucose, and **c)** 5% glucose. The results represent mean values ± standard deviation from three to four independent experiments, including mean values of two technical repetitions. Statistical analyses were performed using one-way ANOVA followed by the Dunnett test vs. control (*\*P* ≤0.05; *\*\*P*≤0.01; *\*\*\*P*≤0.001).

### ADP/ATP ratio variation with quercetin supplementation

Depolarization of the mitochondrial membrane might also affect ATP synthesis, which could augment ADP/ATP ratio. To further determine the influence of quercetin mitochondrial membrane depolarization on oxidative phosphorylation, the ADP/ATP ratio was determined. At 0.1% and 0.5% glucose, quercetin supplementation (0.1 μM and 100 μM) increased the ADP/ATP ratio (**Fig. 2a-b**). In contrast, in cells cultivated with 2% glucose, quercetin supplementation (0.1 μM and 100 μM) did not alter ADP/ATP ratio (**Fig. 2c**). Quercetin at low dose (0.1 μM) only increased ADP/ATP ratio when using 5% glucose (**Fig. 2d**). An increase in the ADP/ATP ratio indicates decreased ATP levels, then our results evidence that quercetin supplementation decreased ATP levels, especially at 0.1% and 0.5% glucose concentrations.

**Fig. 2.**
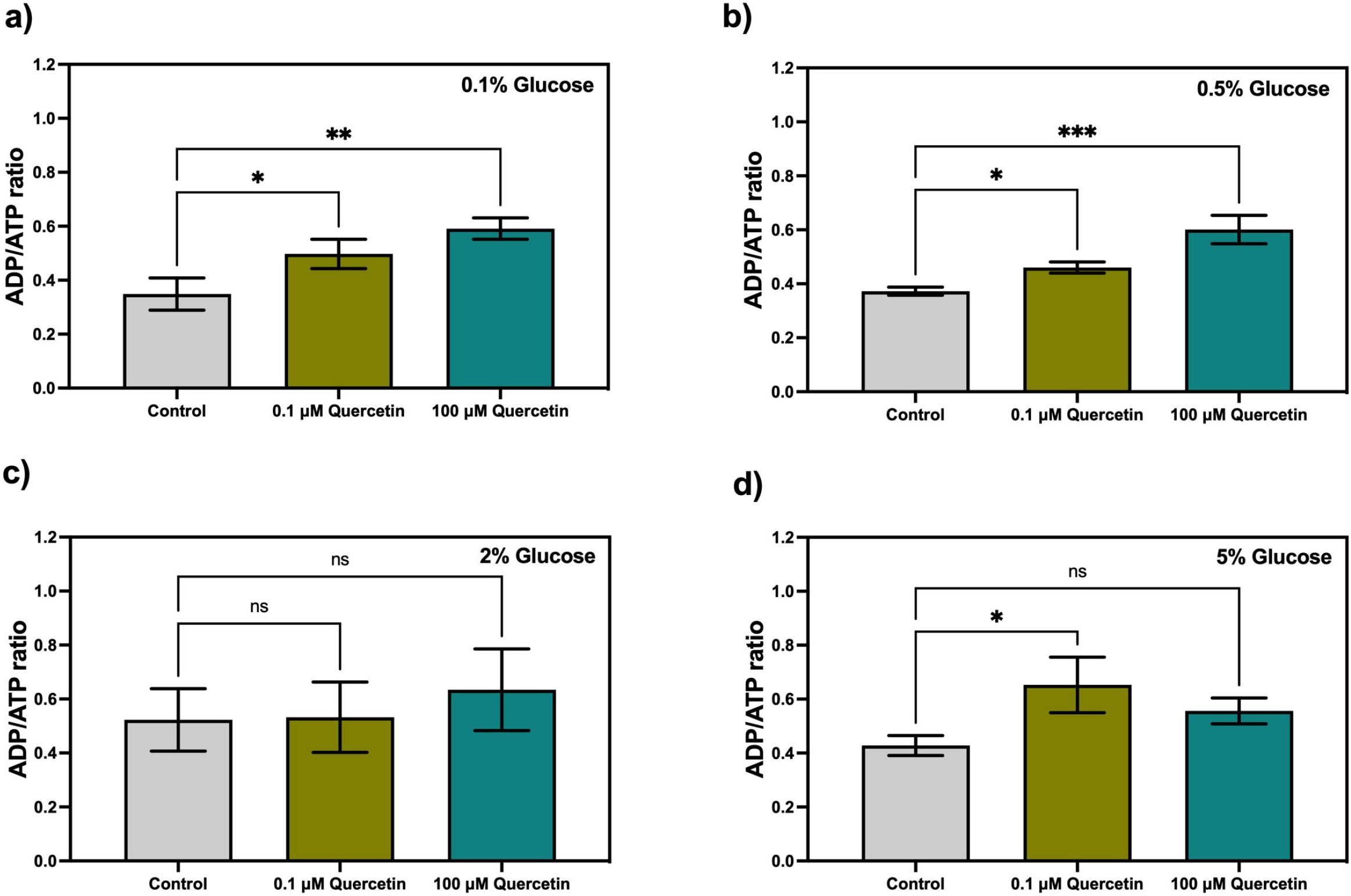
ADP/ATP ratio variation by quercetin supplementation in *S. cerevisiae*. ADP/ATP ratio was measured by luminescence in mid-exponential cultures grown with SC media supplemented with **a)** 0.1% glucose; **b)** 0.5% glucose; **c)** 2% glucose, and **d)** 5% glucose. The results represent mean values ± standard deviation from three independent experiments. Statistical analyses were performed using one-way ANOVA followed by the Dunnett test vs. control (*\*P ≤*0.05; *\*\*P≤*0.01; *\*\*\*P≤*0.001; ns, not-significant).

### Influence of quercetin in ROS generation dependent on glucose concentration

Previously, our research group provided evidence that quercetin supplementation affects mitochondrial respiration and respiratory chain complexes in a glucose and quercetin dose-dependent manner [4]. Herein, we have added evidence that quercetin negatively affects mitochondrial membrane potential and ADP/ATP ratio, indicating that quercetin impairs the oxidative phosphorylation activity. However, the question remains about the fate of electrons missing on the ETC. Quercetin redox potential could promote electrons reduction by taking them from the ETC or also elicit electrons leakage by affecting the fluidization of mitochondrial membranes, while both mechanisms could influence free radical production in cells. We first measured total free radical levels by using the unspecific probe fluorescein to evaluate this idea. Unexpectedly, at 0.1% glucose quercetin supplementation reduced total cellular ROS (**Fig. 3a**), whereas using other glucose levels (0.5%, 2%, and 5%) quercetin did not modify the ROS cellular levels (**Fig. 3b-d**).

**Fig. 3.**
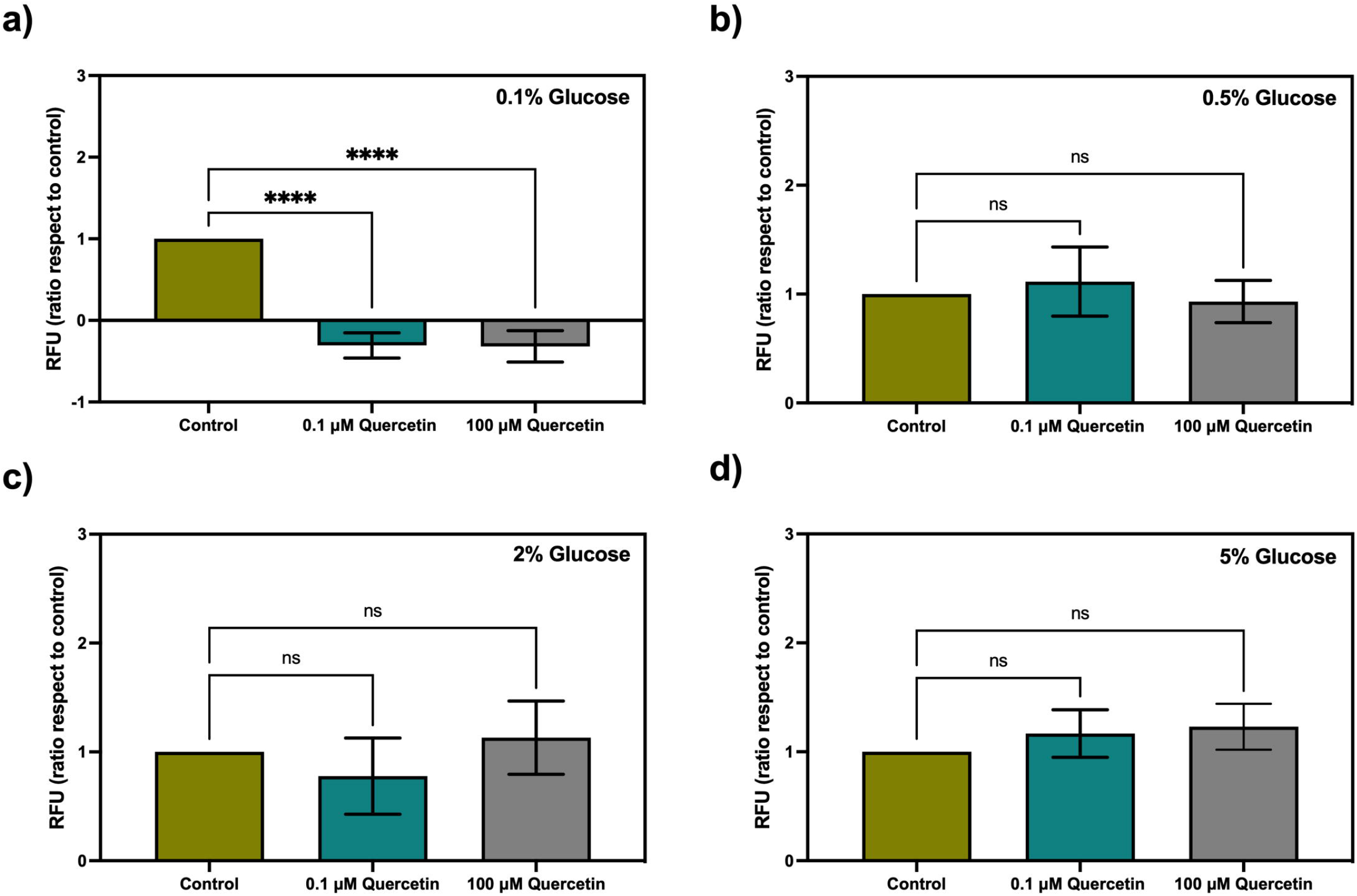
Quantification of total cellular ROS by fluorescein in *S. cerevisiae* cultures supplemented with quercetin. ROS quantification was carried out in mid-exponential cultures grown with SC media supplemented with **a)** 0.1% glucose; **b)** 0.5% glucose; **c)** 2% glucose, and **d)** 5% glucose. The results represent mean values ± standard deviation from five independent experiments, including mean values of two technical repetitions. Statistical analyses were performed using one-way ANOVA followed by the Dunnett test vs. control (*\*\*\*\*P≤*0.0001; ns, not-significant). RFU, relative fluoresce units.

To gain an understanding of quercetin role on ROS levels, we decided to quantify them by MitoSOX, a specific mitochondrial probe for the superoxide anion. Interestingly, quercetin supplementation at a low dose (0.1 μM) increased superoxide anion levels for all glucose concentrations tested (**Fig. 4a-d**). Conversely, 100 μM quercetin only increased superoxide anion levels using 0.1% glucose (**Fig. 4a**). From these apparently contradictory results, the increased production of superoxide anion may have induced the antioxidant cellular response leading to reduced total ROS levels. More experiments are needed in support of this possibility.

**Fig. 4.**
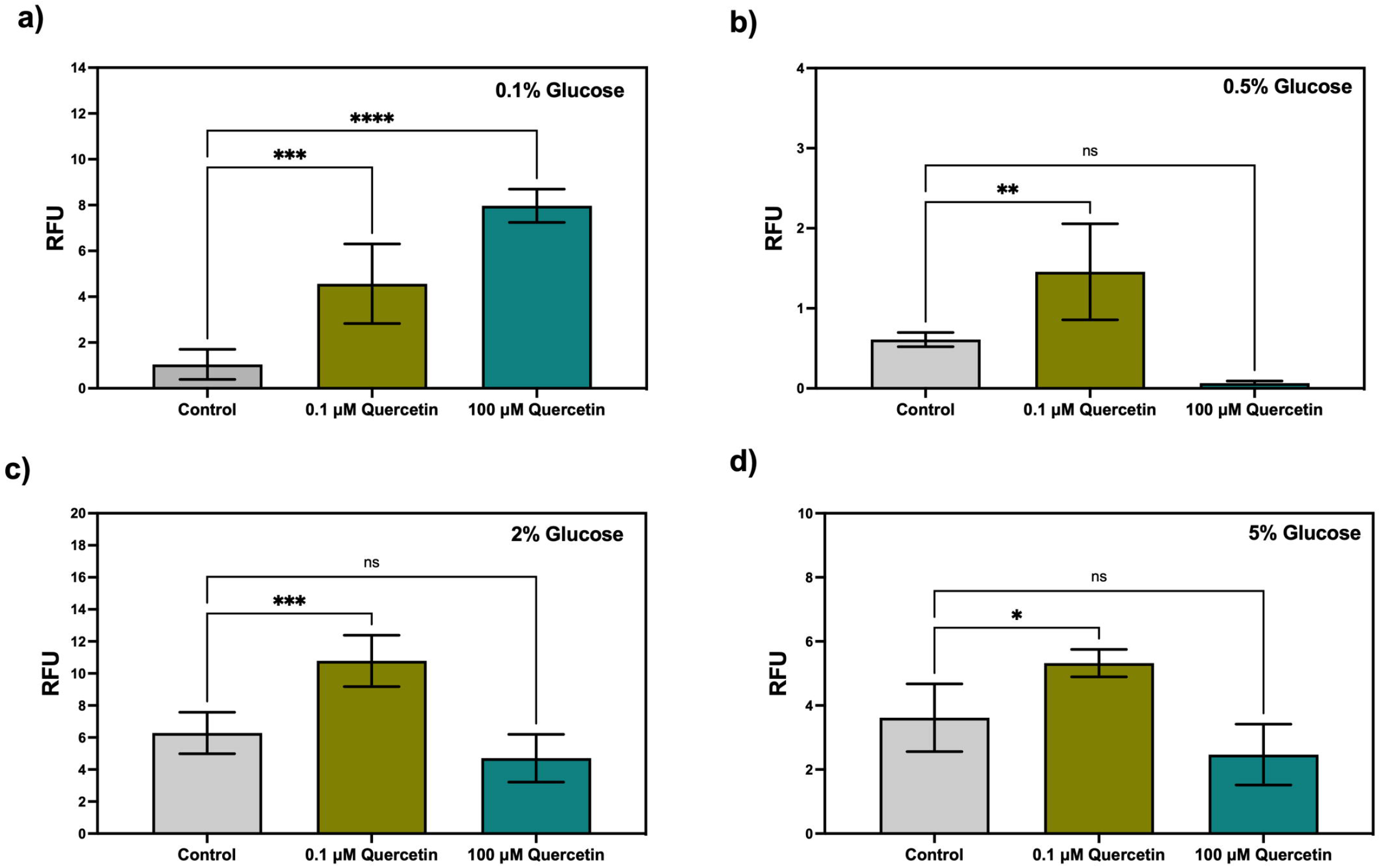
Effect of quercetin supplementation on anion superoxide levels in *S. cerevisiae*. Anion superoxide levels were quantified by fluorescence in mid-exponential cultures grown using SC media supplemented with **a)** 0.1% glucose; **b)** 0.5% glucose; **c)** 2% glucose, and **d)** 5% glucose. The results represent mean values ± standard deviation from five independent experiments, including mean values of two technical repetitions. Statistical analyses were performed using one-way ANOVA followed by the Dunnett test vs. control (*\*P≤*0.05; *\*\*P≤*0.01; *\*\*\*P≤*0.001; *\*\*\*\*P≤*0.0001; ns, not-significant). RFU, relative fluoresce units.

### Influence of quercetin on H_2_O_2_ antioxidant cellular response

To unravel whether quercetin supplementation induces an antioxidant cellular response, and the molecule *per se* is not responsible for the antioxidant phenotype, we performed an assay in which cells were pre-incubated with quercetin followed by washing, and challenged with hydrogen peroxide (H_2_O_2_). This experimental design allowed us to identify the long-standing antioxidant cellular response exerted by quercetin. Two concentrations of H_2_O_2_ (1 mM and 10 mM) reported to exert changes in response to oxidative stress phenotype in *S. cerevisiae*, were used [11, 12]. WT strain cultures responded differently to H_2_O_2_ challenge, at 0.5% glucose, specific growth rate augmented using both H_2_O_2_ levels, while at 5% glucose, sustained growth was only observed using 10 mM H_2_O_2_ (**Fig. 5a**). Pre-incubation with quercetin changed the response patterns facing the oxidative challenge of the WT strain, 0.1 μM and 100 μM quercetin nullified the growth using 1 mM H_2_O_2_ at 0.5% glucose (**Fig. 5a**). Using 5% glucose, 0.1 μM quercetin modified the specific growth rate of the cultures challenged with 1 mM and 10 mM H_2_O_2_ (**Fig. 5b**). However, to better understand whether the change of growth rate patterns exerted by quercetin pre-incubation is related to cellular antioxidant response, we decided to evaluate this phenotype in five strains whose genes codifying for antioxidant-related proteins were blocked. As expected, these mutant strains reverted the phenotype of increased specific growth rate exerted by H_2_O_2_ challenge, observed in the WT strain at both glucose concentrations (**Fig. 6 a-j)**. Importantly, 0.1 μM quercetin pre-incubation restored the growth of *yap1*Δ cultures grown at 5% glucose supplemented with 10 mM H_2_O_2_ (**Fig. 6h**). Interestingly, *sod2*Δ significantly decreased (*P*<0.05) the specific growth rate with 0.1 μM quercetin pre-incubation when was challenged with 10 mM H_2_O_2_ at 0.5% glucose (**Fig. 6e**). This phenotype was not observed using 100 μM quercetin pre-incubation at the same conditions (**Fig. 6e**), and this result coincides with the anion superoxide levels evaluated with MitoSOX that was only increased by 0.1 μM quercetin (**Fig. 4b**). In general, the five mutant strains showed changes in the response to H_2_O_2_, depending on pre-incubation quercetin concentration, and glucose levels (**Fig. 6 a-j**). These data suggest that quercetin induces a long-standing response depending on the antioxidant system.

**Fig. 5.**
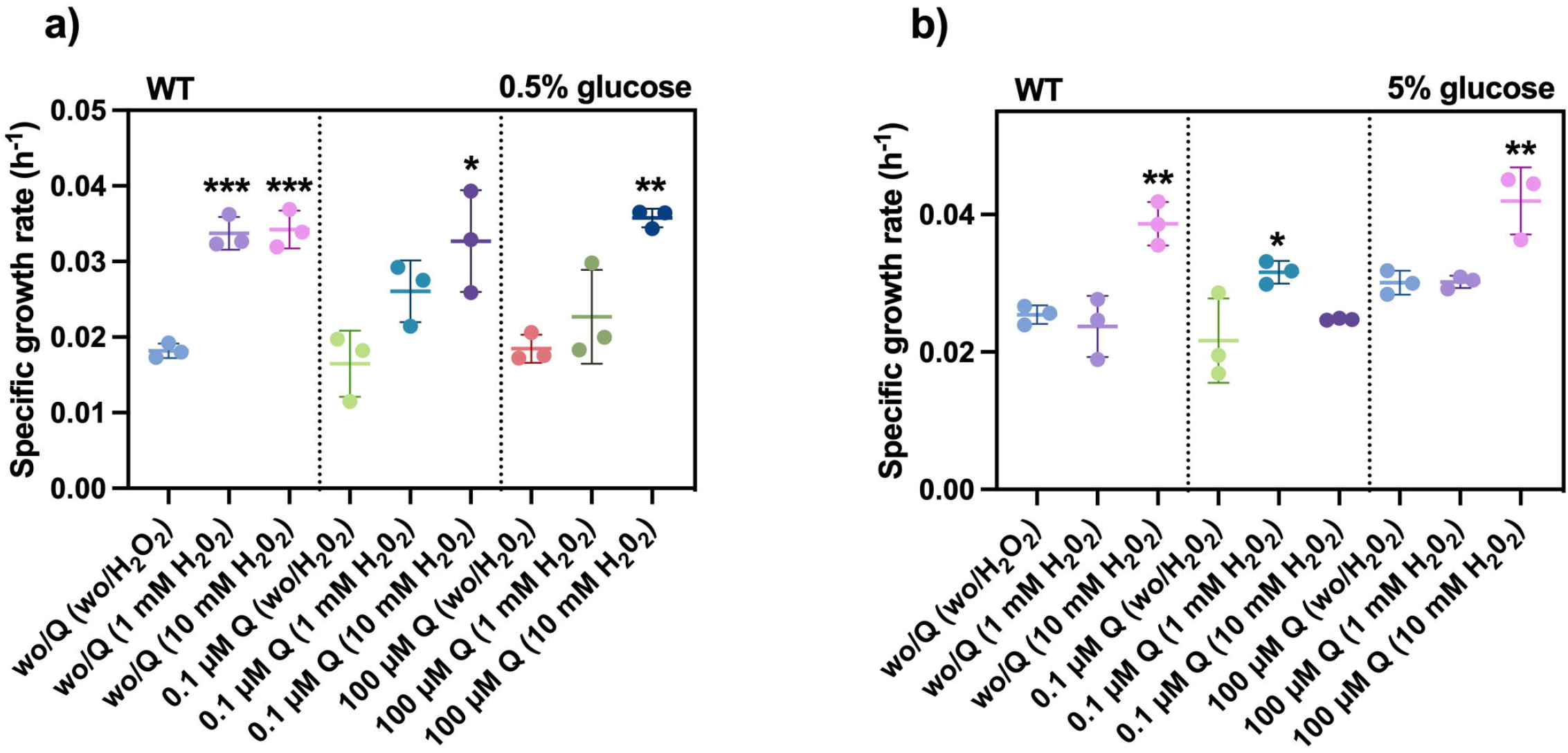
H_2_O_2_ challenge in *S. cerevisiae* cultures pre-incubated with quercetin. WT strain mid-log cultures pre-incubated with (0.1 μM or 100 μM) or without quercetin (Q) were challenged with (1 mM or 10 mM) or without H_2_O_2_ and the exponential growth was measured as a response using the specific growth rate as a parameter, at two glucose concentrations: **a)** 0.5% glucose, and **b)** 5% glucose. The results represent mean values ± standard deviation from three independent experiments, including mean values of three technical repetitions. Statistical analyses were performed using one-way ANOVA followed by the Dunnett test vs. wo/Q (wo/H_2_O_2_); 0.1 μM Q (wo/H_2_O_2_); 100 μM Q (wo/H_2_O_2_), respectively (*\*P≤*0.05; *\*\*P≤*0.01; *\*\*\*P≤*0.001). Q, quercetin; wo, without.

**Fig. 6.**
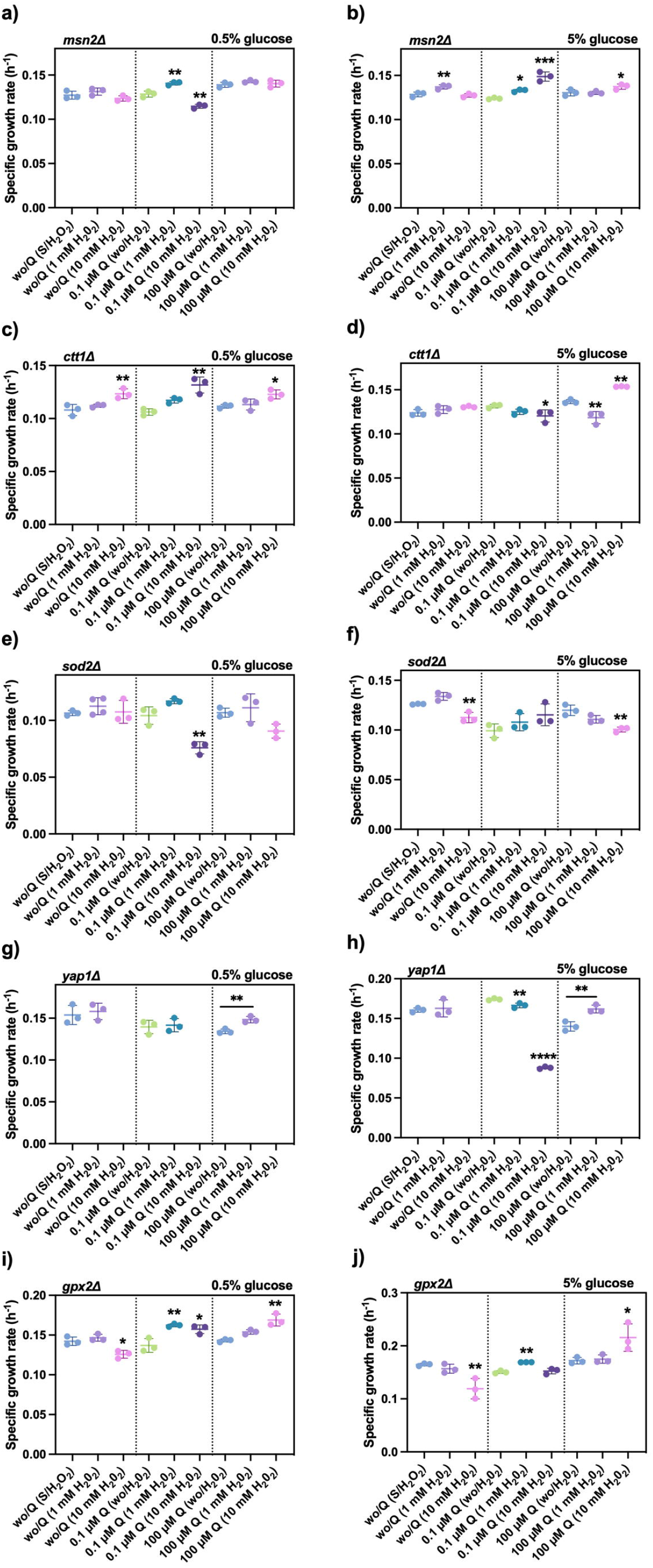
Effect of antioxidant related gene deletant strains cultures pre-incubated with quercetin in growth with H_2_O_2_ challenge. Mid-log cultures of deletant strains pre-incubated with (0.1 μM or 100 μM) or without quercetin (Q) were challenged with (1 mM or 10 mM) or without H_2_O_2_ and the exponential growth was measured as a response using the specific growth rate as a parameter, at two glucose concentrations. **a)** 0.5% glucose, *msn2*Δ; **b)** 5% glucose, *msn2*Δ; **c)** 0.5% glucose, *ctt1*Δ; **d)** 5% glucose, *ctt1*Δ; **e)** 0.5% glucose, *sod2*Δ; **f)** 5% glucose, *sod2*Δ; **g)** 0.5% glucose, *yap1*Δ; **h)** 5% glucose, *yap1*Δ; **i)** 0.5% glucose, *gpx2*Δ; **j)** 5% glucose, *gpx2*Δ. The results represent mean values ± standard deviation from three independent experiments, including mean values of three technical repetitions. Statistical analyses were performed using one-way ANOVA followed by the Dunnett test vs. wo/Q (wo/H_2_O_2_); 0.1 μM Q (wo/H_2_O_2_); 100 μM Q (wo/H_2_O_2_), respectively (*\*P≤*0.05; *\*\*P≤*0.01; *\*\*\*P≤*0.001). Q, quercetin; wo, without.

### Effect of quercetin supplementation on lipoperoxidation

Our hypothesis suggests that quercetin unpairs electron flux in the ETC, probably taking electrons from the ubiquinone-Complex III segment [4], which would result in a quercetin-radical molecule. Because of the hydrophobic nature of quercetin, we propose that quercetin-radical molecules could induce peroxidation of membrane lipids, increasing cellular lipoperoxidation. To test this idea, we performed an experimental design using fatty acids with varying degrees of saturation, and ferric sulfate to induce lipoperoxidation. *S. cerevisiae* incorporates fatty acids from culture media into its membranes, and polyunsaturated fatty acids are more susceptible to peroxidation than saturated fatty acids [13]. For this reason, *S. cerevisiae* cultures were supplemented with three fatty acids with different saturation levels: palmitic 16:0, oleic 18:1, and linoleic 18:2, using two glucose concentrations (0.1% and 5%). Under respiratory growth condition (0.1 % glucose), linolenic acid and ferric sulfate cultures reduced the specific growth rate of *S. cerevisiae* when supplemented with quercetin (0.1 μM and 100 μM) (**Fig. 7c**). Oleic acid supplemented cultures only decreased the specific growth rate at 100 μM quercetin (**Fig. 7b**) and no effect was observed when using palmitic acid and ferric sulfate supplemented cultures in the presence of quercetin (**Fig. 7a**). Under fermentative growth (5% glucose) a reduced growth rate was observed in cultures supplemented with linolenic acid, ferric sulfate and 100 μM quercetin (**Fig. 8c**). Importantly, a fatty acid saturation level-dependent effect was observed under the quercetin influence, upon exponential growth at respiratory growth condition (0.1% glucose), supporting our idea that lipids could suffer peroxidation induced by quercetin. Additionally, these data also show that palmitic acid supplementation increases the quercetin effect on *S. cerevisiae* growth at both glucose concentrations (**Fig. 7a and 8a**).

**Fig. 7.**
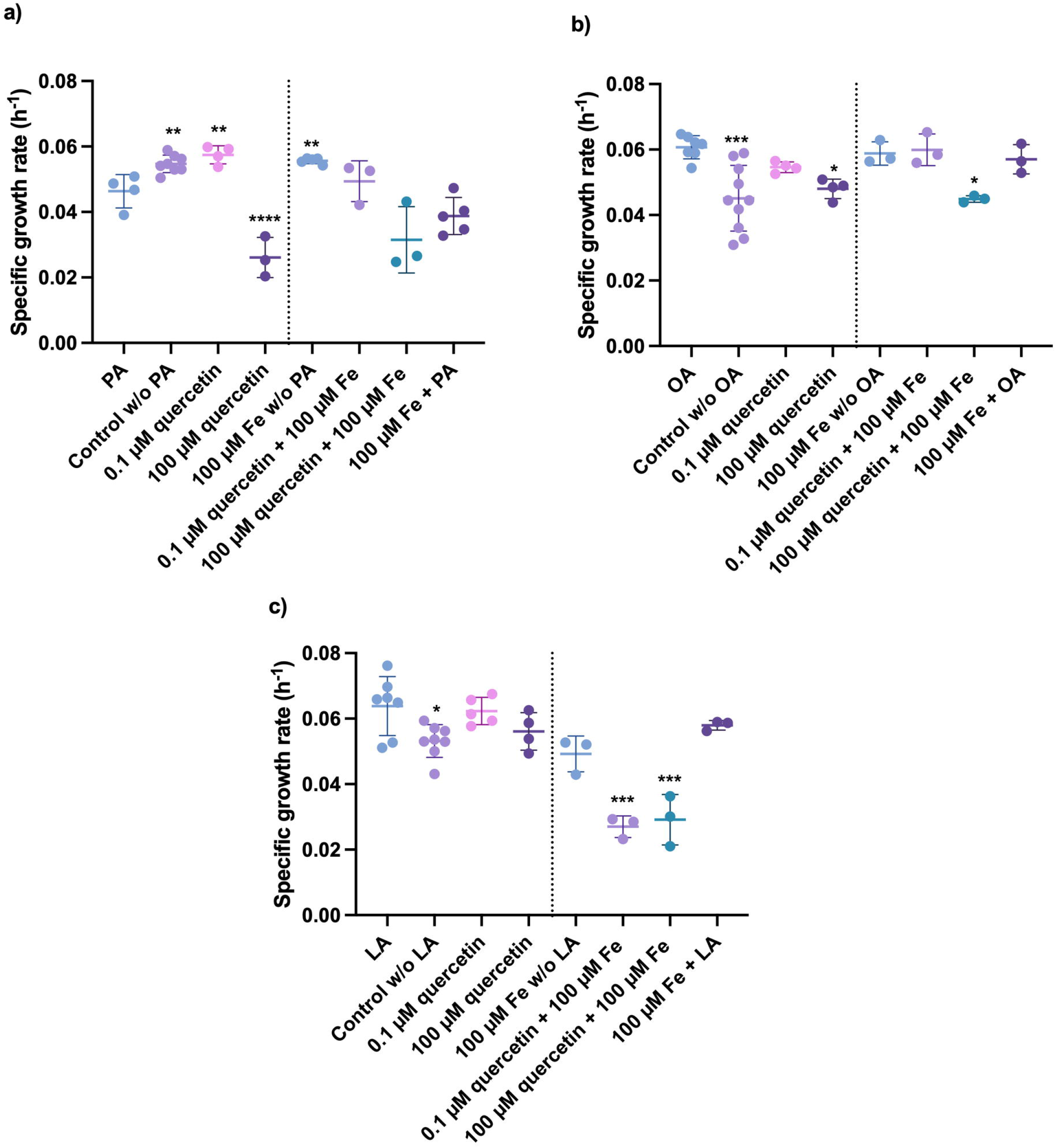
Influence of quercetin on growth of *S. cerevisiae* under lipoperoxidation conditions with different fatty acids. WT cultures at mid-log phase supplemented with quercetin (0.1 μM or 100 μM) at 0.1% glucose with or without 100 μM Fe were assayed to measure its exponential growth using the specific growth rate as a parameter, with different fatty acids. **a)** Palmitic acid (PA); **b)** oleic acid (OA), and **c)** linoleic acid (LA). The results represent mean values ± standard deviation from three to ten independent experiments, including mean values of three technical repetitions. Statistical analyses were performed using one-way ANOVA followed by the Dunnett test vs. PA; OA; LA, respectively (*\*P≤*0.05; *\*\*P≤*0.01; *\*\*\*P≤*0.001; *\*\*\*\*P≤*0.001). wo, without.

**Fig. 8.**
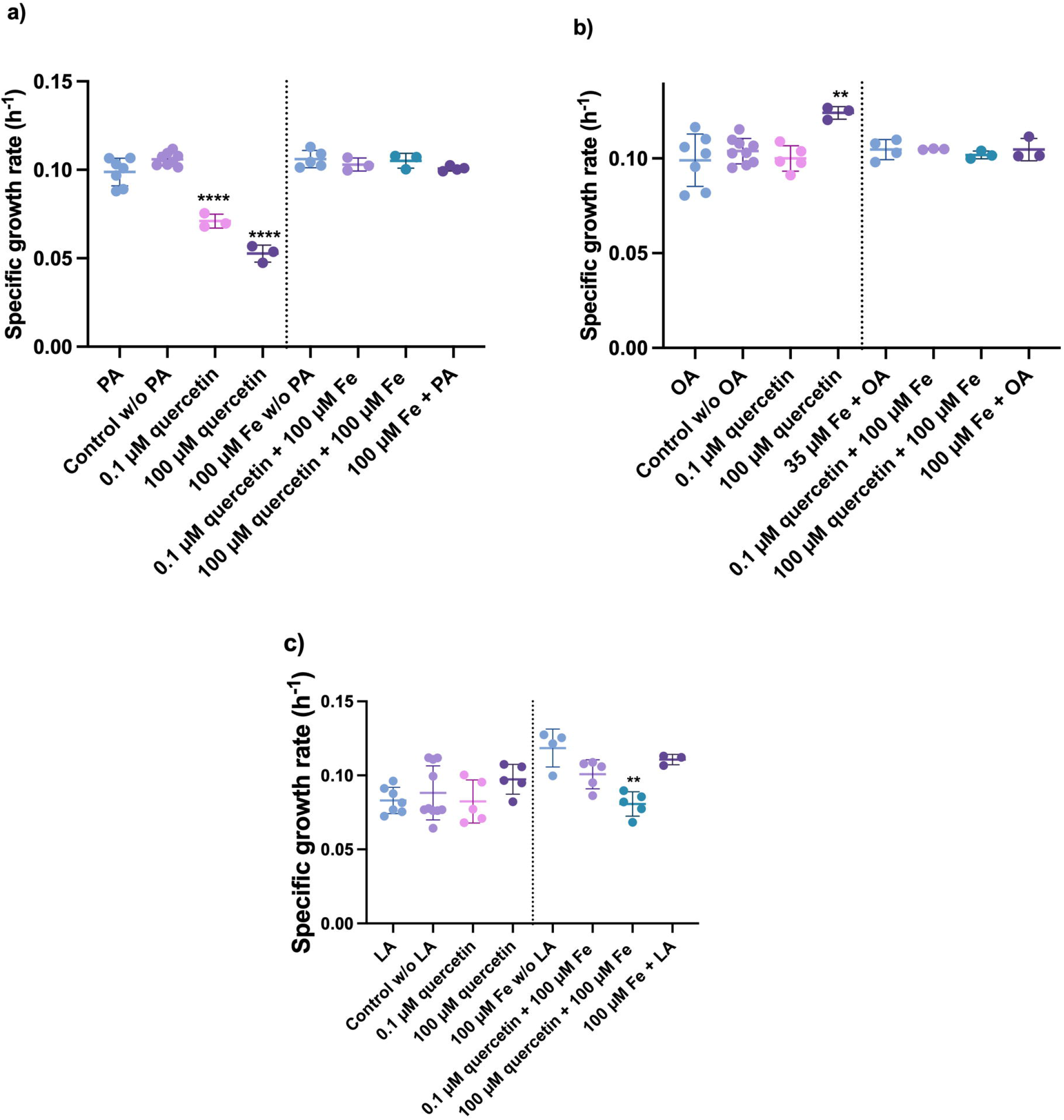
Influence of quercetin on growth of *S. cerevisiae* under lipoperoxidation conditions with different fatty acids. WT cultures at mid-log phase supplemented with quercetin (0.1 μM or 100 μM) at 5% glucose with or without 100 μM Fe were assayed to measure its exponential growth using the specific growth rate as a parameter, with different fatty acids. **a)** Palmitic acid (PA); **b)** oleic acid (OA), and **c)** linoleic acid (LA). The results represent mean values ± standard deviation from three to ten independent experiments, including mean values of three technical repetitions. Statistical analyses were performed using one-way ANOVA followed by the Dunnett test vs. PA; OA; LA, respectively (*\*\*P≤*0.01; *\*\*\*\*P≤*0.001). wo, without.

### Influence of quercetin in cardiolipin levels

Cardiolipin is a key lipid present in the inner mitochondria membrane that is susceptible to lipoperoxidation damage, and if quercetin induces lipoperoxidation it is highly likely that cardiolipin levels are also affected. High quercetin concentration (100 μM) increased cardiolipin levels at the two glucose concentrations tested (**Fig. 9**). Nonetheless, under low quercetin level (0.1 μM), cardiolipin levels were not affected (**Fig. 9**). The increase in cardiolipin levels could explain some differential phenotypes observed at low and high doses of quercetin. However, more evidence is needed to understand the causes of cardiolipin levels increase.

**Fig. 9.**
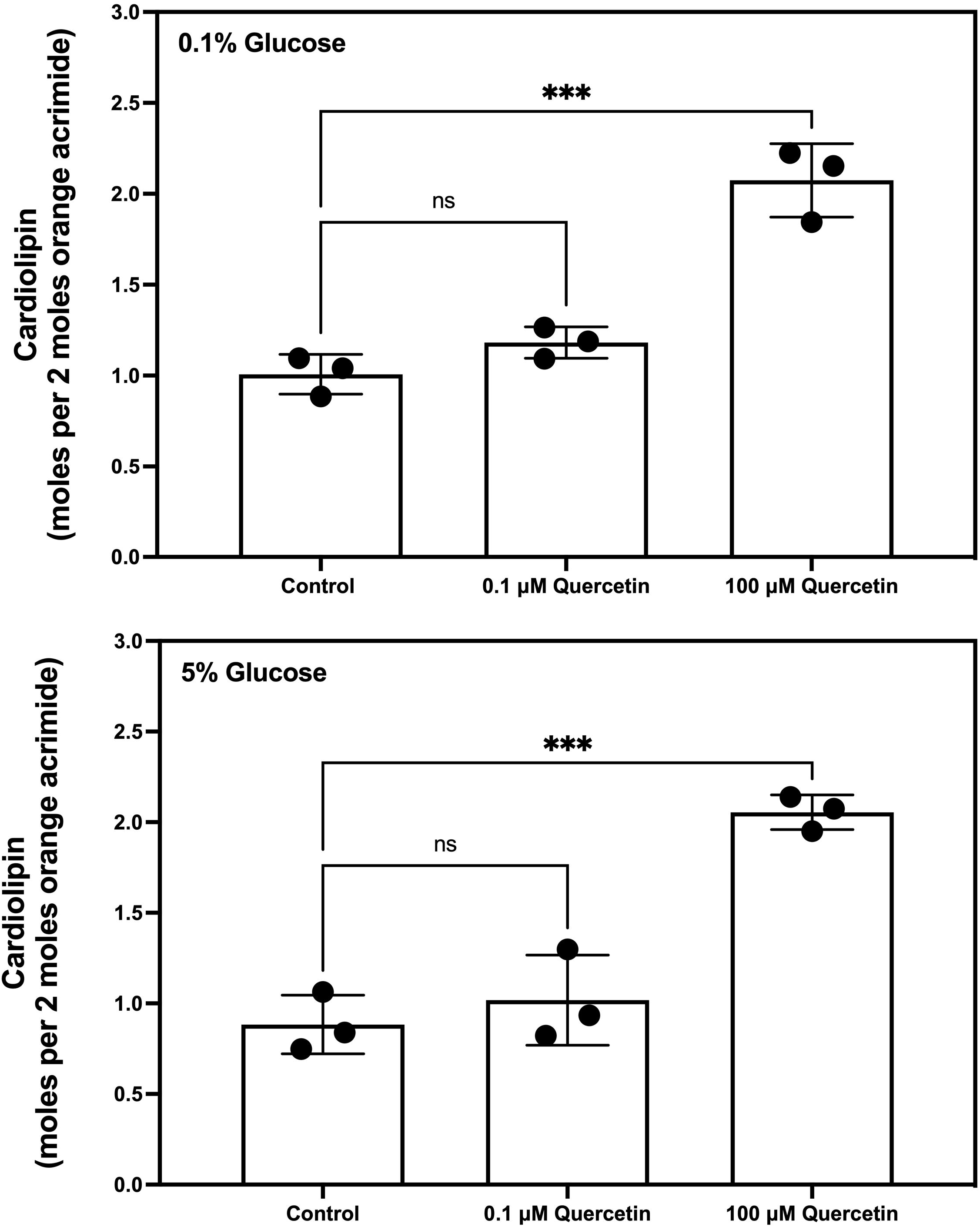
Influence of quercetin supplementation on cardiolipin levels in *S. cerevisiae*. Cardiolipin levels were measured in isolated mitochondria using acridine orange technique from mid-log *S. cerevisiae* cultures supplemented with two quercetin concentrations (0.1 μM or 100 μM) at two glucose concentrations: **a)** 0.1% glucose, and **b)** 5% glucose. The results represent mean values ± standard deviation from three to four independent experiments, including mean values of two technical repetitions. Statistical analyses were performed using one-way ANOVA followed by the Dunnett test vs. control *\*\*\*P≤*0.001; ns, non-significant).

## Discussion

The flavonoid quercetin has emerged as a potential biomedical molecule due to its effect on normalizing risk factors of some chronic non-transmissible diseases [14]. However, the molecular mechanism of quercetin is still not fully understood. Recently, it has been found that quercetin affects electron flux in the ETC at ubiquinone and cytochrome *c* levels, harming mitochondrial respiration [4]. Electron unpairing of quercetin could form free-radical quercetin molecules, which in turn could induce oxidative damage or oxidative stress on cells. The cellular antioxidant system is induced as a response to the oxidative damage caused by free-radical quercetin molecules. Thus, this study aimed to evaluate whether quercetin induces oxidative damage in *S. cerevisiae* cells. Herein, we provided evidence that quercetin supplementation 1) depolarizes the mitochondrial membrane and increases ADP/ATP ratio; 2) increases anion superoxide cellular levels; 3) changes the cellular response to H_2_O_2_ challenge associated to antioxidant cellular response; 4) sensitizes the cellular response to lipoperoxidation challenge, and 4) augments cardiolipin levels; all the phenotypes are dependent on quercetin and glucose concentrations. Overall, these data indicate quercetin induces a pro-oxidant effect related to mitochondrial dysfunction.

Oxidative phosphorylation is the main target of quercetin [7]. The redox potential of quercetin is related to its capacity for electron flux disruption on the ETC. Accordingly, quercetin reduces the mitochondrial respiration of *S. cerevisiae* in a glucose and quercetin-dependent manner [4]. In this study, we proved that quercetin also depolarizes the mitochondrial membrane (**Fig. 1**). The impairing of mitochondrial respiration and depolarization of the mitochondrial membrane by quercetin could also affect ATP production. For example, in mitochondria isolated from male Wistar rats, quercetin supplementation (25-50 μM) decreased ATP levels [15]. Moreover, in submitochondrial particles of beef heart (*Bos primigenius taurus*), quercetin supplementation (4, 12, and 20 μg/mL) inhibited mitochondrial ATPase [16]. Herein, we found that quercetin supplementation (0.1 μM and 100 μM) increased the ADP/ATP ratio in cells cultured using 0.5% glucose **(Fig. 2)**. Using 5% glucose, only 0.1 μM quercetin supplementation increased the ADP/ATP ratio. The increase of the ADP/ATP ratio, evidence the reduction in ATP levels, suggesting that quercetin reduction pf ATP levels is dependent on oxidative phosphorylation disturbance.

Our main idea is based on the fact that quercetin could take electrons from the ETC, suggesting that it might become a free radical molecule or induce free radical formation, but data supporting this idea is still lacking. Here, we found that low quercetin level (0.1 μM) increased the anion superoxide levels (**Fig. 3**) in all glucose concentrations tested, whereas high dose (100 μM) only increased anion superoxide at 0.1% glucose (**Fig. 3a**). Induction of superoxide production by 50 μM quercetin was also reported in colon cancer cells (HCT116) [5]. Also, mitochondriotropic quercetin derivative 7-O-(4-triphenylphosphoniumbutyl) quercetin iodide (5 μM) acts as a pro-oxidant augmenting superoxide anion production in mitochondria [17]. Induction of anion superoxide production has also been related to changes in the mitochondrial permeability transition pore (MPTP), whose onset is associated with high quercetin dose supplementation (20-50 μM) [5]. The opening of MPTP could be related to the oxidative damage induced by quercetin, whereas the opening of MPTP by ROS protects mitochondria from oxidative stress [18].

The pro-oxidant effect of quercetin might induce cellular antioxidant systems. This idea suggests that quercetin at the cellular level does not cause the antioxidant phenotype *per se*; instead, it causes a pro-oxidant effect that elicits an antioxidant system. In support of this possibility, we found that the H_2_O_2_ challenge in quercetin pre-incubated cells changes growth patterns in WT (**Fig. 5**) and in antioxidant deletant strains (**Fig. 6**). Free radical quercetin-derived molecules could promote lipoperoxidation or exacerbate lipoperoxidation conditions since quercetin is a highly hydrophobic molecule. As expected, quercetin enhances the toxic effect of molecules that induce lipoperoxidation of the fatty acids tested in this work, in a glucose and quercetin concentration-dependent manner (**Fig. 7 and 8**). Interestingly, quercetin supplementation did not decrease cardiolipin mitochondrial content, on the contrary, a high dose (100 μM) of quercetin increased cardiolipin content (**Fig. 9**).

Overall this results indicate that quercetin promotes a pro-oxidant phenotype associated with mitochondrial dysfunction that induce the cellular antioxidants systems.

## Acknowledgments

This study was funded by grants from Tecnológico Nacional de México (17716.23-PD) and PICIR 2022 program from the Instituto de Ciencia, Tecnología e Innovación del Estado de Michoacán de Ocampo (Grant: PICIR-008). The authors would like to thank M.C Adriana González-Gallardo and Dra. Anaid Antaramian (Unidad de Proteogenómica del INB-UNAM) for their technical support.

## Data availability statement

The data that support the findings of this study are available from the corresponding author upon reasonable request.

## Author contribution

All authors contributed to the study conception and design. Material preparation, data collection, and analysis were performed by Andres Carrillo-Garmendia, Ana Leticia Vaca-Martinez and Blanca Lucia Carmona-Moreno. The first draft of the manuscript was written by Luis Alberto Madrigal-Perez and Carlos Regalado-Gonzalez, and all authors commented on previous versions of the manuscript. All authors read and approved the final manuscript.

